# PKProbDesign: RNA inverse folding including pseudoknots by optimizing thermodynamic folding probability

**DOI:** 10.64898/2026.07.09.736945

**Authors:** Takumi Otagaki, Junichi Iwakiri, Goro Terai, Kiyoshi Asai, Kengo Sato

**Affiliations:** School of Life Science and Technology, Institute of Science Tokyo, Japan; Department of Computational Biology and Medical Sciences (CBMS), The University of Tokyo, 5-1-5 Kashiwanoha, Kashiwa, Chiba, 277-8561, Japan; AI-Empowered Life Science Initiative (ALIS), the Joint Support-Center for Data Science Research (DS), Research Organization of Information and Systems (ROIS), 4-1-1 Toranomon, 105-6923, Tokyo, Japan

## Abstract

**Motivation:** RNA inverse folding, the design of RNA sequences that fold into specified target secondary structures, is a central problem in RNA design, with applications in functional RNA engineering, synthetic biology, and nucleic-acid therapeutics. This task becomes especially challenging for pseudoknotted target structures because pseudoknots break the nested structure assumed by standard thermodynamic folding models. Existing pseudoknot inverse-folding methods often rely on structure-predictor-based objectives. These methods do not directly optimize the probability that a sequence folds into the specified pseudoknotted target structure. Such optimization requires an evaluator that can assign a folding probability to the specified target within a pseudoknot-aware ensemble.

**Results:** We present PKProbDesign, a sampling-based inverse-folding framework that directly optimizes a thermodynamic folding-probability objective for pseudoknotted targets. For each target, sampled sequences are scored by combining the folding probability of a pseudoknot-free scaffold with the conditional folding probability of the remaining extension component. On 254 unique density-2 PseudoBase++ targets, PKProbDesign achieved the highest pseudo-joint folding probability on 245 targets, compared with 6 for DesiRNA, 3 for MODENA, and none for antaRNA.

**Conclusions:** PKProbDesign demonstrates that pseudoknot inverse folding can be formulated with target folding probability as its objective rather than structure-prediction agreement alone. By combining scaffold decomposition with conditional folding-probability evaluation based on CParty, the method provides a practical folding-probability-based approach to designing sequences for density-2 pseudoknotted targets.

**Availability:** The source code of PKProbDesign is available at https://github.com/TakumiOtagaki/PKProbDesign.

## 1. Introduction

RNA inverse folding is the problem of designing an RNA sequence that folds into a given target secondary structure. It is a fundamental task in functional RNA engineering and nucleic-acid design [Gao et al., 2010, Kleinkauf et al., 2015b, Tang et al., 2026, Zhou et al., 2026]. Classical inverse-folding methods have often evaluated candidate sequences by comparing their minimum-free-energy (MFE) structures with the target structure [Dotu et al., 2014, Busch and Backofen, 2006].

However, the MFE structure alone does not describe the thermodynamic ensemble of possible secondary structures associated with a sequence. Consequently, even when the MFE structure matches the target, the target does not necessarily dominate this ensemble, because the ensemble can contain many alternative structures with similar free energies [McCaskill, 1990]. This observation has motivated ensemble-aware design objectives such as target folding probability. Target folding probability is the probability assigned to the target structure within this ensemble [Zadeh et al., 2011, Zhou et al., 2023, Tang et al., 2026, Zhou et al., 2026].

A recent example of this probability-based approach is SamplingDesign [Tang et al., 2026]. It reformulates RNA design as a continuous optimization problem over a probability distribution on valid sequences, rather than as a search over individual sequences. By approximating the target folding-probability objective and its gradient via Monte Carlo sampling, it achieved strong performance on pseudoknot-free targets. This formulation represents the valid sequence space through continuous distributional parameters, making gradient-based optimization possible.

However, because this folding-probability objective is defined for pseudoknot-free secondary structures, the formulation does not directly address pseudoknotted targets. Extending this probability-based strategy to pseudoknots requires an evaluator that can assign target-specific probabilities within a pseudoknot-aware thermodynamic ensemble.

RNA pseudoknots are structural motifs that play important roles in many functional RNAs [Pleij et al., 1985, Brierley et al., 2007]. However, allowing pseudoknots breaks the nested structure assumed by pseudoknot-free secondary-structure models. Without restricting the structural class, exact pseudoknotted RNA secondary-structure prediction is NP-hard [Akutsu, 2000, Lyngsø and Pedersen, 2000].

Several methods have been developed for pseudoknot inverse folding, including antaRNA [Kleinkauf et al., 2015a], MODENA [Taneda, 2011], and DesiRNA [Wirecki et al., 2025]. antaRNA scores each sequence using the distance between the structure predicted by pKiss [Theis et al., 2010] and the target, together with GC-content and sequence constraints. MODENA uses IPknot [Sato et al., 2011] or HotKnots [Ren et al., 2005] to predict each structure. It optimizes two objectives: the similarity of the predicted structure to the target and its free energy. DesiRNA optimizes the folding probability of the first pseudoknot-free layer. For the remaining pseudoknot pairs, it repeatedly predicts MFE structures and ranks its outputs by how closely the resulting structure matches the target. It therefore does not evaluate the folding probability of the complete pseudoknotted target. None of these methods directly optimizes a folding-probability objective that includes all target base pairs, including those that form pseudoknots.

HFold [Jabbari et al., 2007] and CParty [Gray et al., 2024] define a hierarchical folding framework for density-2 structures, a restricted class defined by constraints on crossing base pairs [Jabbari et al., 2008]. HFold takes a pseudoknot-free scaffold as input to predict a density-2 pseudoknotted structure. It finds the set of additional base pairs that gives the lowest free energy when added to the scaffold. CParty instead computes the partition function over all possible sets of additional base pairs for the same scaffold. This conditional partition function enables the computation of conditional folding probabilities under the density-2 thermodynamic model. Together, HFold and CParty provide the probability calculations needed to score specified targets in this class.

Here, we present PKProbDesign (pseudoknot folding probability-based design), a sampling-based inverse-folding framework for density-2 pseudoknotted targets. PKProbDesign decomposes each target into a pseudoknot-free scaffold and an extension component. PKProbDesign scores sampled sequences by combining the folding probability of the scaffold with the conditional folding probability of the extension component. It then optimizes the sequence distribution to increase this combined probability using gradient estimates obtained from sampling. The scaffold folding probability is calculated with LinearPartition [Zhang et al., 2020], and the conditional extension probability is calculated with CParty.

## 2. Materials and Methods

### 2.1 Overview of PKProbDesign

PKProbDesign divides each density-2 pseudoknotted target into a pseudoknot-free scaffold and an extension component, combines the scaffold folding probability and conditional extension folding probability, and optimizes a sequence distribution by sampling. Figure 1 summarizes these steps from dividing the target to selecting the final sequences. At each iteration, PKProbDesign samples RNA sequences from the distribution, evaluates their combined probabilities, and updates the distribution parameters using gradient estimates obtained from these samples.

**Fig. 1.**
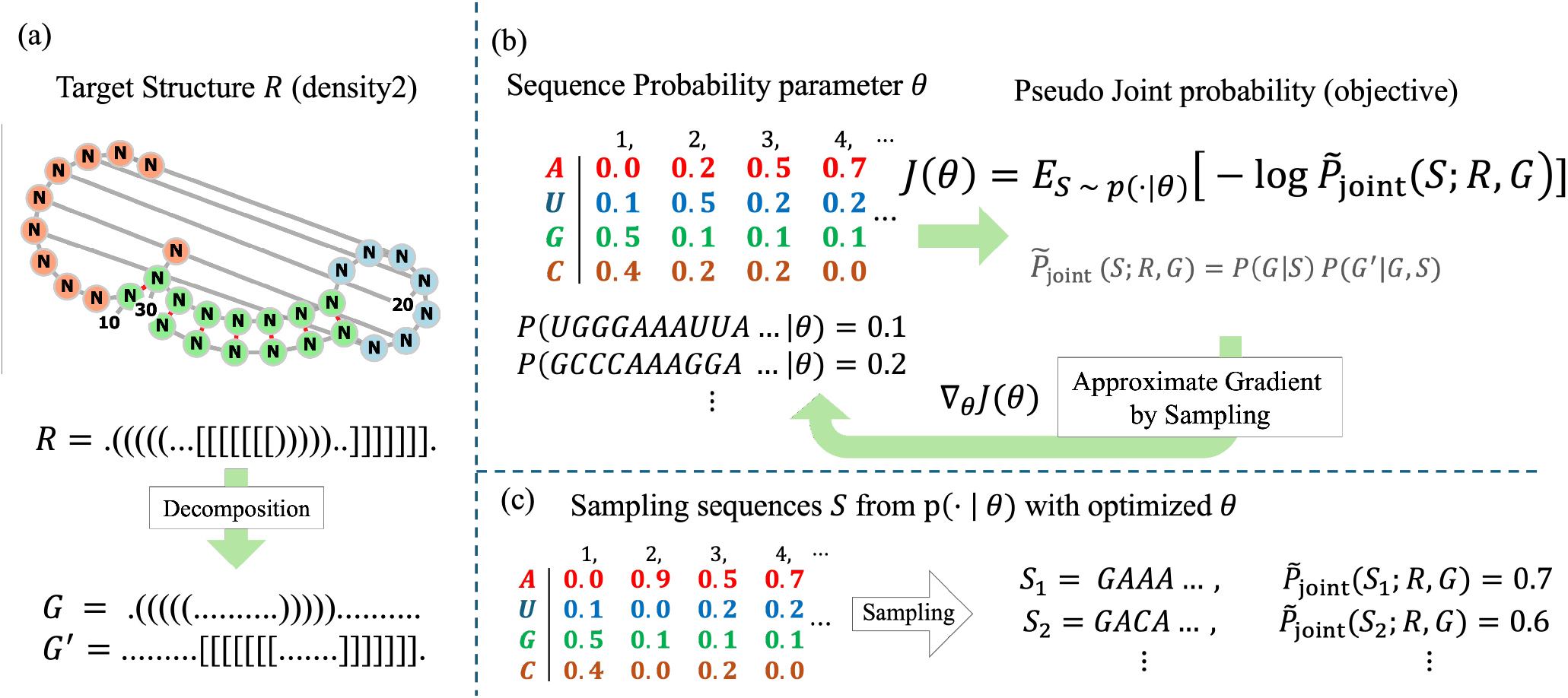
Overview of PKProbDesign. (a) A density-2 target structure *R* is divided into two pseudoknot-free components, a scaffold *G* and an extension component *G*′, with *R* = *G ∪ G*′. (b) Sequences *S* are sampled from the distribution *p*_*θ*_(*S*), whose parameters are *θ*. The pseudo-joint folding probability is 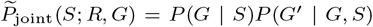, the product of the scaffold folding probability and conditional extension folding probability. The objective *J* (*θ*) is the expected negative logarithm of this probability under *p*_*θ*_(*S*), and its gradient ∇_*θ*_*J* (*θ*) is approximated using the sampled sequences. (c) Sequences retained during optimization are combined with additional sequences sampled from the optimized *p*_*θ*_(*S*) and ranked by the pseudo-joint folding probability 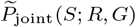. Higher values indicate higher folding probabilities.

### 2.2. Terminology

Throughout this paper, an RNA secondary structure is represented as a set of base pairs (*i, j*) with *i < j*, where *i* and *j* are sequence positions. Accordingly, *R* denotes the base-pair set of a target pseudoknotted secondary structure, *S* an RNA sequence, *G* a pseudoknot-free scaffold base-pair set, and *G*′ a pseudoknot-free extension-component base-pair set. When a target is decomposed into these two parts, we write *R* = *G* ∪ *G*′ with *G* ∩ *G*′ = ∅; in other words, *G* and *G*′ are disjoint base-pair sets. This decomposition is not necessarily unique. The deterministic crossing-graph procedure in Section 2.4 divides *R* into two pseudoknot-free components, *G*_big_ and *G*_small_. It assigns as many target base pairs as possible to *G*_big_ and the remaining base pairs to *G*_small_.

We follow the density-2 definition of Jabbari et al. [2008]. In this definition, base pairs involved in crossing interactions are organized into bands. Base pairs within a band are nested, meaning that one base pair encloses another. If a base pair outside a band crosses a base pair in the band, it crosses all base pairs in that band. For each pseudoknot, density is the largest number of its bands that span the same sequence position. A structure is density-2 if every pseudoknot has density at most two.

### 2.3. Sampling-based optimization of the sequence distribution

Following Tang et al. [2026], we optimize a probability distribution *p*_*θ*_(*S*) over the valid sequence space instead of optimizing one sequence directly. Each unpaired position is represented by a 4-way categorical variable over {*A, C, G, U*}. For each base pair (*i, j*), the two nucleotides are represented jointly by a 6-way categorical variable over canonical base pairs {*AU, UA, GC, CG, GU, UG*}, so that the probability of assigning bases at *i* and *j* is modeled as a joint distribution over the pair rather than as two independent single-base marginals. Thus, every sampled sequence satisfies the canonical-pair (Watson-Crick and Wobble) constraints at all positions paired in the target

We minimize the expected score under *p*_*θ*_(*S*):

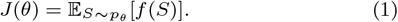

The function *f* (*S*) is treated as a sequence score; its definition for density-2 pseudoknotted targets is given in Section 2.4. Differentiating the expectation in Eq. 1 with respect to *θ* and using ∇_*θ*_*p*_*θ*_(*S*) = *p*_*θ*_(*S*)*∇*_*θ*_ log *p*_*θ*_(*S*), we obtain

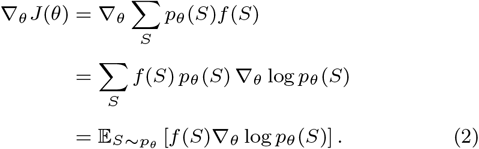

In practice, at each iteration we sample *K* = 30 sequences *S*_1_, …, *S*_*K*_ ~ *p*_*θ*_ and approximate the gradient by

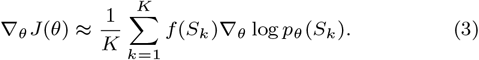

Thus, this gradient-based update does not require a differentiable internal evaluator *f* (*S*).

The sequence-distribution parameters for the position-wise and pair-wise categorical variables are represented as logits and transformed into probabilities by a softmax. At each iteration, we sample *K* sequences from *p*_*θ*_(*S*) and evaluate *f* (*S*_*k*_) for each sample. We then construct the gradients and update the parameters using Adam [Kingma and Ba, 2014], including the standard first- and second-moment bias corrections, for 100 iterations. We retain sequences sampled and scored during optimization, sample 30 additional sequences from the optimized distribution, and retain the 20 sequences with the highest pseudo-joint folding probabilities as the design pool.

### 2.4. Target decomposition and pseudo-joint folding probability

Before optimization, we construct one scaffold-extension decomposition for each density-2 target structure from its base-pair crossing graph. Each vertex in this graph represents a target base pair, and two vertices are connected when their base pairs cross. For a density-2 target, this graph is bipartite. Dividing its base pairs into two pseudoknot-free components is therefore a graph-colouring problem with two colours: vertices joined by an edge must be assigned to different groups. For each component, we assign the group with more base pairs to *G*_big_ and the other group to *G*_small_. If the two groups contain the same number of base pairs, the assignment is determined from their base-pair positions. Applying this rule independently to every connected component gives *G*_big_ the largest possible number of base pairs while keeping both components pseudoknot-free. Together, *G*_big_ and *G*_small_ contain all base pairs in *R*. The implementation checks every pair of the *N* target base pairs when constructing the graph, requiring *O*(*N* ^2^) time. In the main analysis, we set *G* = *G*_big_ and *G*′ = *G*_small_. We also evaluate the reversed scaffold assignment as a supplementary analysis, with *G* = *G*_small_ and *G*′ = *G*_big_

For a decomposed target *R* = *G ∪ G*′, we define a pseudo-joint folding probability for that decomposition,

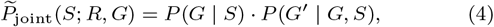

where *P* (*G* | *S*) is the scaffold folding probability. Following SamplingDesign [Tang et al., 2026], we evaluate this probability with LinearPartition [Zhang et al., 2020]. No beam pruning occurred in the calculations reported here. The term *P* (*G*′ | *G, S*) is the conditional folding probability of the extension component given the scaffold; its definition is given in Section 2.5.

The optimization score is the negative logarithm of this probability:

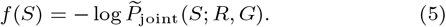

Let *D*(*R*) denote the set of pairs (*G, G*′) such that *G* and *G*′ are pseudoknot-free, *G ∩ G*′ = *∅*, and *G ∪ G*′ = *R*. Under the hierarchical folding assumption, summing over all valid decompositions would give the full joint probability under this model, *P*_joint_(*S*; *R*):

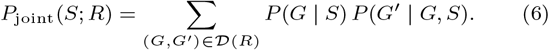

Each term corresponds to one possible scaffold-extension decomposition of the target. Because every term in the sum is nonnegative, the pseudo-joint probability for the selected decomposition is a lower bound on *P*_joint_(*S*; *R*). However, to keep the computation tractable, we optimize only the term that uses *G*_big_ as the scaffold, 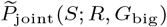, rather than evaluating the full sum.

### 2.5. CParty-based conditional-probability evaluation

We consider the class of density-2 pseudoknotted structures handled by HFold and CParty. All CParty-based conditional-probability and extension-prediction calculations use the DP09 energy parameters [Andronescu et al., 2010]. HFold predicts the minimum-free-energy density-2 structure that extends a given pseudoknot-free scaffold *G*. CParty computes the corresponding partition function over the extensions allowed by the hierarchical folding model. For a sequence *S* and scaffold *G*, we define

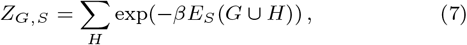

where β is the inverse temperature constant, *E*_*S*_(·) denotes the free energy of the specified structure for sequence *S*, and the sum ranges over all pseudoknot-free structures *H* such that *G ∩ H* = *∅* and *G* ∪ *H* is density-2. The associated conditional probability is given by

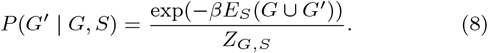

This probability measures the conditional folding probability of the extension component on top of the given scaffold, with the Boltzmann weight evaluated for the full union structure *G ∪ G*′, not for *G*′ alone. We use *P* (*G*′ | *G, S*) as the inverse-folding score for the specified target extension.

### 2.6. Structure-prediction protocols and evaluation metric

We considered two protocols that obtain scaffolds from each designed sequence:

- **Two-stage prediction.**We first predict one pseudoknot-free scaffold from the designed sequence with LinearFold using beam size 100, and then predict the minimum-free-energy extension conditionally on that scaffold.
- **HotKnots-guided prediction.**HotKnots uses stable stems to explore alternative pseudoknotted structures [Ren et al., 2005]. We use up to 20 HotKnots-derived scaffold candidates and also include the empty scaffold. For each scaffold, we predict the minimum-free-energy extension and retain the resulting full structure with the lowest energy.

Neither protocol uses distance to the target when selecting a predicted structure. For both protocols, we compared the selected structures with the target using normalized base-pair Hamming distance. If *P*_pred_ and *P*_tgt_ denote the predicted and target base-pair sets, respectively, and *P* [*i, j*] denotes the indicator of whether base pair (*i, j*) is present in a set *P*, then the distance is

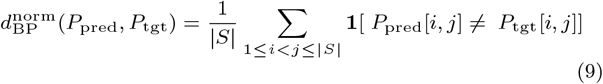

where |S| is the sequence length and **1**[·] equals 1 when its condition is true and 0 otherwise.

## 3. Results

### 3.1. Benchmark dataset

We constructed the benchmark dataset from PseudoBase++ [Taufer et al., 2009], a database of annotated RNA pseudoknot structures. For each entry, we extracted the RNA sequence and its annotated pseudoknotted secondary structure. In the dot-bracket notation, paired positions are represented by matching bracket symbols and unpaired positions by dots. We retained density-2 targets with sequence length at most 100 nt for which every target base pair (*i, j*) satisfies *j − i −* 1 ≥ 3. Four entries were excluded because structure data were unavailable. These filters yielded 325 entries, which were deduplicated by exact target dot-bracket structure to give 254 unique targets. All 254 targets can be evaluated by the conditional folding model used in this study with *G*_big_ as the scaffold. The target lengths range from 21 to 100 nt, with mean 49.12 nt and median 44 nt; the full distribution is shown in Supplementary Figure S1.

### 3.2. Objective performance and method comparison

We compared PKProbDesign with antaRNA [Kleinkauf et al., 2015a], DesiRNA [Wirecki et al., 2025], and MODENA [Taneda, 2011] using the pseudo-joint folding probability 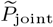 defined in Eq. 4. For every designed sequence, we computed log *P* (*G* | *S*), log *P* (*G*′ | *G, S*), and log 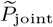 using *G*_big_ as the scaffold and *G*_small_ as the extension. PKProbDesign used this decomposition directly during sequence design. DesiRNA uses () and [] for the two pseudoknot-free components and treats the () component as its first layer. Before running DesiRNA, we reassigned the dot-bracket symbols without changing the target base pairs so that *G*_big_ was encoded by () and *G*_small_ by []. MODENA and antaRNA do not use this scaffold-extension decomposition during design. For every target and method, we evaluated a pool of 20 designed sequences and used the sequence with the highest pseudo-joint folding probability as the representative design.

For PKProbDesign, we combined sequences retained during optimization with 30 additional samples from the optimized distribution and selected the 20 sequences with the highest pseudo-joint folding probabilities. No target-distance measure was used to select representative sequences. The probability comparison evaluates the sequence pools under the folding model used by PKProbDesign; it does not provide a method ranking independent of the evaluation model.

Figure 2 summarizes the probability metrics for the representative sequences. The left panel shows the pseudo-joint folding probabilities, and the other two panels show the scaffold and conditional-extension folding probabilities separately.

**Fig. 2.**
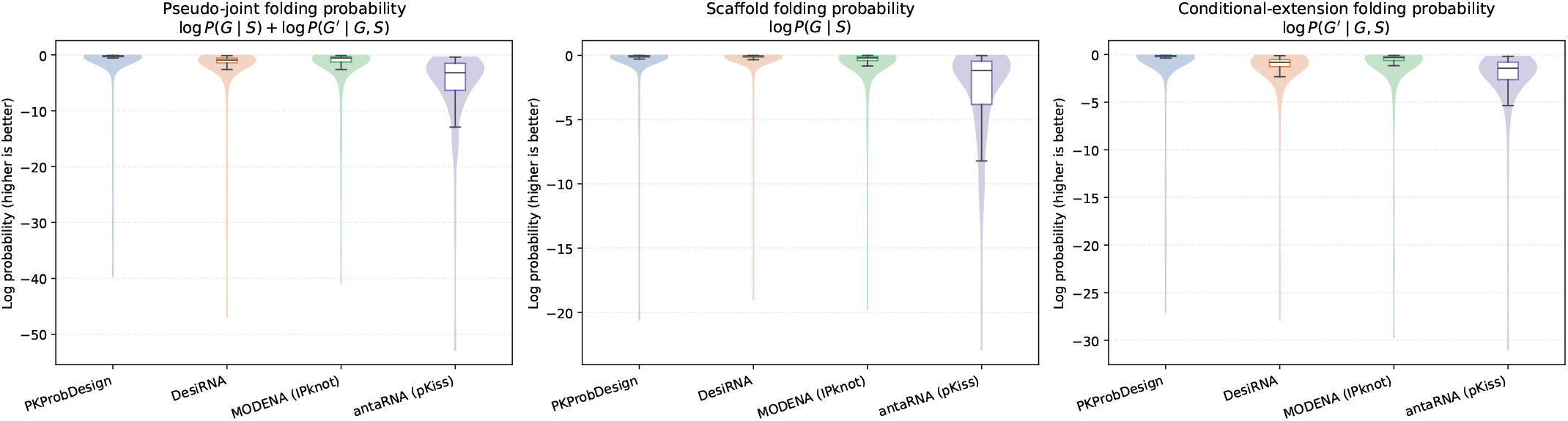
Thermodynamic folding-probability metrics for the final 254-target benchmark. For each target and method, the sequence with the highest pseudo-joint folding probability among 20 designs is shown. In the panel labels, *G* = *G*_big_ is the scaffold and *G*′ = *G*_small_ is the extension for every method. The left, middle, and right panels show the log pseudo-joint folding probability, scaffold folding probability log *P* (*G* | *S*), and conditional extension folding probability log *P* (*G*′ | *G, S*), respectively. Higher values indicate higher folding probabilities.

PKProbDesign achieved the highest pseudo-joint folding probability on 245 of the 254 targets, compared with 6 for DesiRNA, 3 for MODENA, and none for antaRNA. In the paired comparison with DesiRNA, PKProbDesign had the higher pseudo-joint folding probability on 248 targets and the lower value on 6 targets. The median log-probability difference, calculated as PKProbDesign minus DesiRNA, was 0.627 (Bonferroni-adjusted two-sided Wilcoxon signed-rank *p* = 4.91 × 10^*−*39^). The scaffold terms were comparable: PKProbDesign was higher on 126 targets and DesiRNA on 128, with mean log probabilities of −0.291 and −0.257, respectively (unadjusted *p* = 0.777). In contrast, PKProbDesign had the higher conditional-extension term on 252 targets, with mean values of −0.671 for PKProbDesign and *−*1.642 for DesiRNA. Thus, the difference in pseudo-joint folding probability arose primarily from the conditional-extension term rather than from the scaffold term. The full paired results are reported in Supplementary Table S1. In separate PKProbDesign runs, using *G*_big_ as the scaffold gave a higher pseudo-joint probability than using *G*_small_ on 246 of 254 targets; the reversed assignment gave the higher value on the remaining 8 targets. The main comparison used the *G*_big_ assignment fixed by graph colouring before optimization rather than selecting the higher-scoring assignment for each target. Reversing the assignment changed the relative performance of PKProbDesign and DesiRNA (Supplementary Text S3 and Figure S4); the corresponding structure-prediction results are reported in Supplementary Figures S5 and S6.

Run parameters and software versions for the compared methods are listed in Supplementary Texts S1 and S2.

### 3.3. Structure-prediction-based evaluation

We next evaluated the designed sequences by comparing predicted structures with the target using normalized base-pair Hamming distance (Eq. 9). As described in Section 2.6, we used the two-stage and HotKnots-guided prediction protocols, which obtain scaffolds from LinearFold and HotKnots, respectively. Figure 3 summarizes the resulting comparisons.

**Fig. 3.**
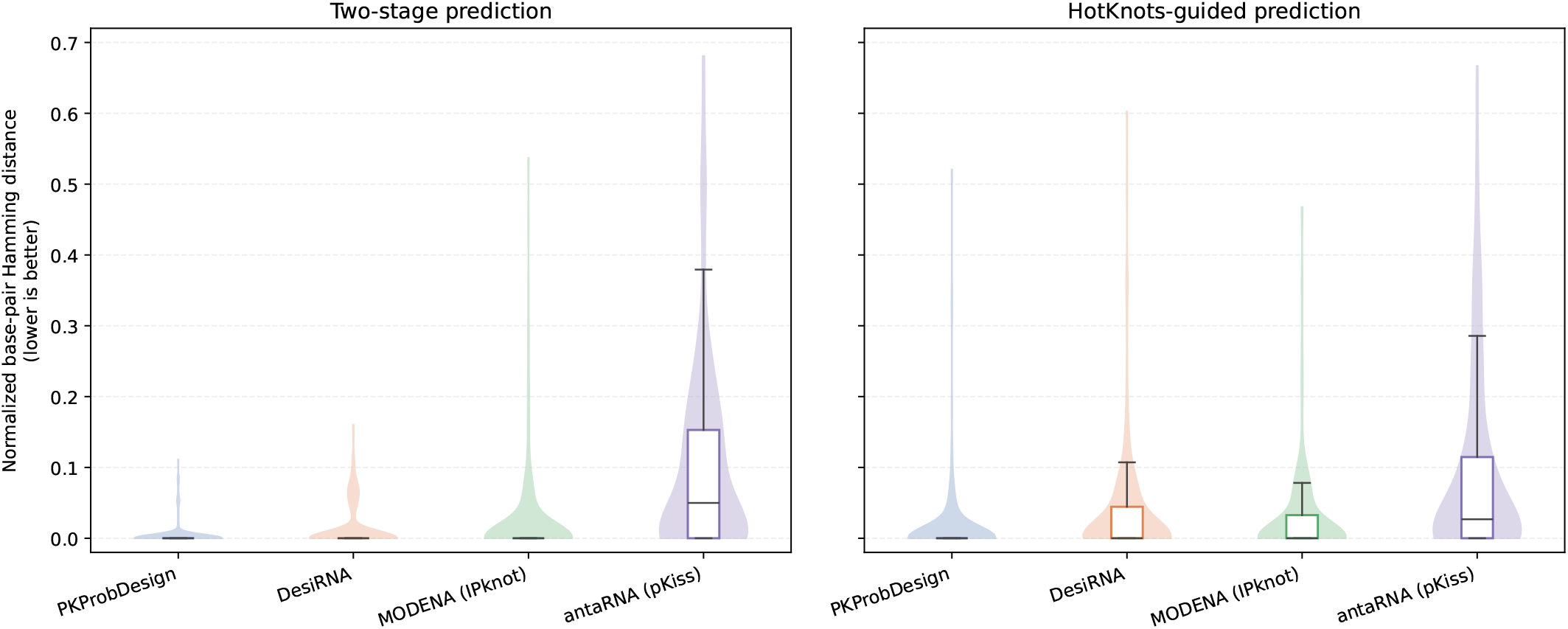
Structure-prediction results for the protocols defined in Section 2.6. The left and right panels show normalized base-pair Hamming distance for two-stage and HotKnots-guided prediction, respectively. Exact-match rates for PKProbDesign, DesiRNA, MODENA, and antaRNA were 92.9%, 79.1%, 79.1%, and 42.9%, respectively, under two-stage prediction, and 81.9%, 65.4%, 70.5%, and 46.1% under HotKnots-guided prediction. For every method, the same representative sequence shown in Figure 2 was used. The scaffolds used for prediction were obtained from LinearFold or HotKnots. Lower distance indicates better agreement with the target structure.

Under two-stage prediction, PKProbDesign had the lowest mean normalized distance and the highest exact-match rate (0.0040; 92.9%), followed by DesiRNA (0.0145; 79.1%), MODENA (0.0214; 79.1%), and antaRNA (0.0993; 42.9%). Under HotKnots-guided prediction, PKProbDesign again had the lowest mean normalized distance and the highest exact-match rate (0.0133; 81.9%), followed by MODENA (0.0280; 70.5%), DesiRNA (0.0330; 65.4%), and antaRNA (0.0875; 46.1%). Supplementary Figure S3 reports the corresponding normalized base-pair Hamming-distance summary obtained from IPknot predictions.

### 3.4. Representative case study

Figure 4 shows PKB00076, a selected success case in which PKProbDesign has high pseudo-joint folding probability and its HotKnots-guided prediction exactly matches the target. MODENA also gives an exact prediction for this target, whereas the representative antaRNA and DesiRNA sequences have raw base-pair Hamming distances of 6 and 4, respectively. Normalization is unnecessary in this within-target comparison because all predictions have the same sequence length. This example illustrates that high pseudo-joint folding probability can coincide with accurate structure prediction, but it is presented as a selected example rather than as a benchmark-wide pattern.

**Fig. 4.**
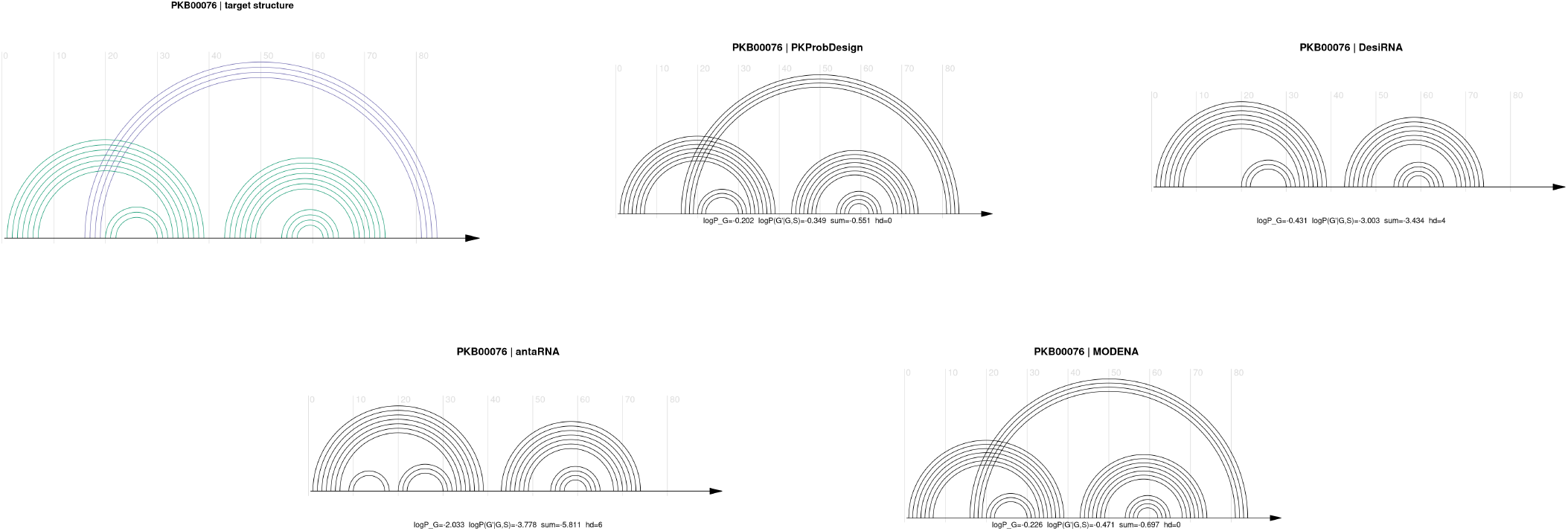
Selected success case for PKB00076 under HotKnots-guided prediction. The top row shows the target structure, the predicted structure for PKProbDesign, and the predicted structure for DesiRNA. The bottom row shows the predicted structures for antaRNA and MODENA. Each designed-sequence panel is annotated with the method name, PKB identifier, log *P* (*G* | *S*), and log *P* (*G*′ | *G, S*). The secondary-structure drawings were rendered with R4RNA/R-chie [Tsybulskyi et al., 2020].

## 4. Discussion

### 4.1 Folding-probability scores and structure-prediction results

In this study, we formulated inverse folding for density-2 pseudoknotted targets by combining target decomposition, conditional folding-probability evaluation, and sampling-based optimization. Across the benchmark, PKProbDesign achieved the highest pseudo-joint folding probability on 245 of 254 targets and had a higher conditional-extension probability than DesiRNA on 252 targets. These results show that target folding probability can be used directly as an optimization objective for density-2 pseudoknot inverse folding.

Pseudo-joint folding probability and structure-prediction agreement capture different outcomes. PKProbDesign directly optimizes pseudo-joint folding probability, which is the primary evaluation criterion in this study. Structure prediction provides a complementary comparison between predicted and target structures.

The larger scaffold leaves fewer target base pairs in the conditional extension, which may help explain the higher pseudo-joint probability observed when *G*_big_ was used as the scaffold. However, this experiment does not show that the smaller extension caused the higher probability.

### 4.2. Scope of the density-2 class

The proposed method is defined for density-2 pseudoknotted targets, and the score-based benchmark in this study is restricted accordingly. The benchmark analyzed here contains 254 unique density-2 PseudoBase++ targets that satisfy the sequence-length and minimum-loop criteria. The results therefore establish performance within this selected target class, but do not imply that density-2 covers all RNA pseudoknots. For longer or structurally more complex RNAs, a broader structural class may become necessary. The present formulation is therefore limited to this target class and does not characterize natural RNA secondary structures in general.

### 4.3. Relationship to existing pseudoknot inverse-folding methods

Existing pseudoknot inverse-folding methods such as antaRNA [Kleinkauf et al., 2015a], DesiRNA [Wirecki et al., 2025], and MODENA [Taneda, 2011] use different search strategies and scoring functions to design sequences for pseudoknotted target structures. In contrast, the proposed approach optimizes a folding probability that includes both components of a specified density-2 pseudoknotted target rather than optimizing predicted-structure agreement itself. The comparisons therefore characterize the methods using both pseudo-joint folding probability and agreement between predicted and target structures.

DesiRNA provides the closest probability-based comparison. Its default Ed-Epf score corresponds to the negative logarithm of the folding probability of the pseudoknot-free input scaffold. For pseudoknotted targets, DesiRNA constructs additional layers through repeated MFE structure prediction after masking positions paired in the preceding layers. Similarity between this multilayer prediction and the target is used to rank the designed sequences, but the default search objective does not include a conditional folding probability for the extension. In the main comparison, which used *G*_big_ as the scaffold, DesiRNA and PKProbDesign had comparable scaffold probabilities, with no significant target-wise difference. PKProbDesign instead had a higher conditional-extension term on 252 of 254 targets. Thus, the improvement in its pseudo-joint score arose primarily from the conditional pseudoknot extension rather than from a failure of DesiRNA to stabilize the scaffold.

MODENA uses IPknot as its structure predictor during optimization. It had the highest exact-match rate under IPknot prediction (65.4%; Supplementary Figure S3) and ranked second under HotKnots-guided prediction (70.5%; Figure 3), but achieved the highest pseudo-joint folding probability on only 3 of 254 targets. The IPknot result is consistent with the predictor used during MODENA optimization, whereas the pseudo-joint comparison measures the probability assigned to the specified scaffold-extension decomposition under the folding model used by PKProbDesign.

antaRNA directly optimizes structural agreement under pKiss together with a specified GC-content target. Its lower pseudo-joint probabilities and larger structure-prediction distances under the evaluation models used here may reflect differences between its optimization objective and the measures used in this study. The GC-content constraint may also affect the search, but its contribution was not evaluated separately.

### 4.4. Methodological limitations

Several limitations of the present study should be noted. First, the main objective is defined for one scaffold-extension decomposition. The full target probability under the hierarchical model would require summing over all valid decompositions (*G, G*′), whereas the main analysis uses the single deterministic assignment with *G*_big_ as the scaffold.

Second, although SamplingDesign [Tang et al., 2026] proposed higher-order coupled variables for terminal mismatches and trimismatches, the present work restricts the sequence distribution to canonical base-paired couplings.

Third, downstream structural and biological validation remain limited in the present study, and the biological significance of the folding-probability objective will require further evaluation.

### 4.5. Future directions

The present study suggests several directions for future work. First, the scaffold folding probability is already evaluated with LinearPartition, whose beam size can be adjusted for longer sequences. A similar linear-time approximation of the CParty-based conditional calculation could reduce the cost of both probability calculations and enable longer targets and larger density-2 benchmarks to be explored.

Second, beyond comparisons based on structure prediction, the present objective should be tested more directly through experimental validation, for example using DMS/PairMap-style probing or in vitro folding assays.

## 5. Acknowledgements

This study was carried out using the TSUBAME4.0 supercomputer at the Institute of Science Tokyo.

