## Supplementary Information for "PKProbDesign: RNA inverse folding including pseudoknots by optimizing thermodynamic folding probability"

### Figure S1

Figure S1 summarizes the sequence-length distribution of the density-2 PseudoBase++ benchmark used in this study.

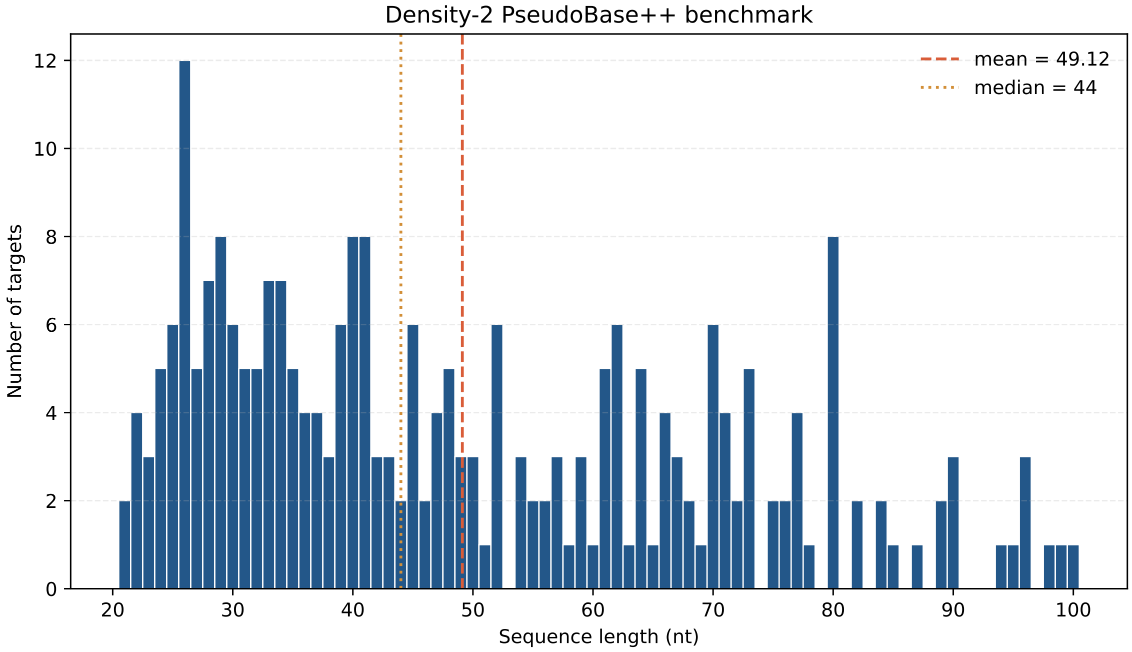


Figure S1: Length distribution of the 254 unique density-2 PseudoBase++ targets used in the main analysis. The mean and median lengths are 49.12 nt and 44 nt, respectively, with a range of 21–100 nt. The 1–40, 41–70, and 71–100 nt intervals contain 112, 95, and 47 targets, respectively.

### Text S1

The benchmark and method-comparison settings were as follows:

- target length at most 100 nt, every target pair $(i, j)$ satisfying $j - i - 1 \geq 3$, and 254 unique density-2 target structures obtained by deduplicating 325 filtered entries;
- 100 optimization iterations with standard Adam bias correction, 30 sampled sequences per iteration, 15 optimizer threads, retention of the top 20 designs, and 30 additional samples from the optimized distribution;
- 20 designs per target and method;
- DesiRNA used its default `Ed-Epf:1.0` score, pseudoknot mode, Turner 1999 parameters, 10 replicas, exchange interval 100, a 60-second budget, and 20 designs; ACGU-content control and negative design were disabled;
- antaRNA used a GC-content soft target of 0.5, a 60-second budget, 20 designs, and pKiss pseudoknot prediction;
- PKProbDesign used $G_{\text{big}}$ as the scaffold and $G_{\text{small}}$ as the extension during design; DesiRNA uses `()` and `[]` to represent two pseudoknot-free target pair classes in dot-bracket notation. Before design, these symbols were assigned without changing the target base-pair set so that $G_{\text{big}}$ was encoded as the `()` input scaffold and $G_{\text{small}}$ as the `[]` second component;
- MODENA and antaRNA do not use this decomposition during design; their resulting designs were evaluated using $G_{\text{big}}$ as the scaffold and $G_{\text{small}}$ as the extension for probability evaluation and representative selection;
- CParty conditional-probability and Viterbi calculations used the DP09 energy parameters;
- LinearPartition was compiled in Vienna mode with `-Dlpv` and used without beam pruning;
- two-stage prediction used LinearFold with beam size 100 followed by conditional Viterbi prediction;
- HotKnots-guided prediction used at most 20 hotspot-derived scaffolds plus the empty scaffold, ran conditional Viterbi prediction for every proposal, and selected the prediction with the lowest energy.

Target distance was not used to select representative sequences, scaffold proposals, or predicted structures.

### Text S2

#### Software versions

The software versions used for the comparison were:

- **PKProbDesign**: version `v1.0.0`.
- **LinearPartition**: `b450fb3`, Vienna mode.
- **DesiRNA**: `v1.0-57-gbdb4908`.
- **MODENA**: version `v0040`.
- **antaRNA**: `v2.0.1.2`, with pKiss and ViennaRNA 2.7.2.
- **LinearFold**: SamplingDesign implementation, revision `e88f883f`.
- **HotKnots**: version 2.0, revision `b01a2c9`.
- **IPknot**: version 1.1.0.

### Table S1

Table S1 reports paired comparisons of PKProbDesign and DesiRNA using $G_{\text{big}}$ as the scaffold and $G_{\text{small}}$ as the extension. Positive differences favor PKProbDesign. Raw two-sided Wilcoxon values and Bonferroni-adjusted values for the three displayed probability metrics are reported.

Table S1: Paired comparison of PKProbDesign and DesiRNA on 254 unique density-2 targets using $G_{\text{big}}$ as the scaffold. The paired difference is defined as PKProbDesign minus DesiRNA.

| Metric | $n$ | PKProbDesign better | DesiRNA better | Ties | Median $\Delta$ | Raw $p$ | Adjusted $p$ |
|---|---|---|---|---|---|---|---|
| $\log \widetilde{P}_{\text{joint}}$ | 254 | 248 | 6 | 0 | 0.6266 | $8.18 \times 10^{-40}$ | $4.91 \times 10^{-39}$ |
| $\log P(G \mid S)$ | 254 | 126 | 128 | 0 | $-0.0008$ | $7.77 \times 10^{-1}$ | 1.00 |
| $\log P(G' \mid G, S)$ | 254 | 252 | 2 | 0 | 0.6127 | $2.84 \times 10^{-42}$ | $1.71 \times 10^{-41}$ |

### Figure S2

Figure S2 compares separate PKProbDesign runs using $G_{\text{big}}$ or $G_{\text{small}}$ as the design scaffold. Let $S_{\text{big}}$ and $S_{\text{small}}$ denote the representative sequences from the corresponding runs. We define the probability difference shown in the left panel as

$$\begin{aligned}\Delta_{\text{prob}} = \; & \log P(G_{\text{big}} \mid S_{\text{big}}) + \log P(G_{\text{small}} \mid G_{\text{big}}, S_{\text{big}}) \\ & - \log P(G_{\text{small}} \mid S_{\text{small}}) - \log P(G_{\text{big}} \mid G_{\text{small}}, S_{\text{small}}).\end{aligned}$$

For either the two-stage or HotKnots-guided protocol $q$, let $\widehat{R}_q(S)$ denote the structure predicted from $S$. We define the corresponding distance difference as

$$\Delta_{\text{dist}}^{(q)} = d_{\text{BP}}^{\text{norm}}\left(\widehat{R}_q(S_{\text{small}}), R\right) - d_{\text{BP}}^{\text{norm}}\left(\widehat{R}_q(S_{\text{big}}), R\right).$$

Here, $d_{\text{BP}}^{\text{norm}}$ is the normalized base-pair Hamming distance. Positive values of either difference favor the $G_{\text{big}}$-scaffold run.

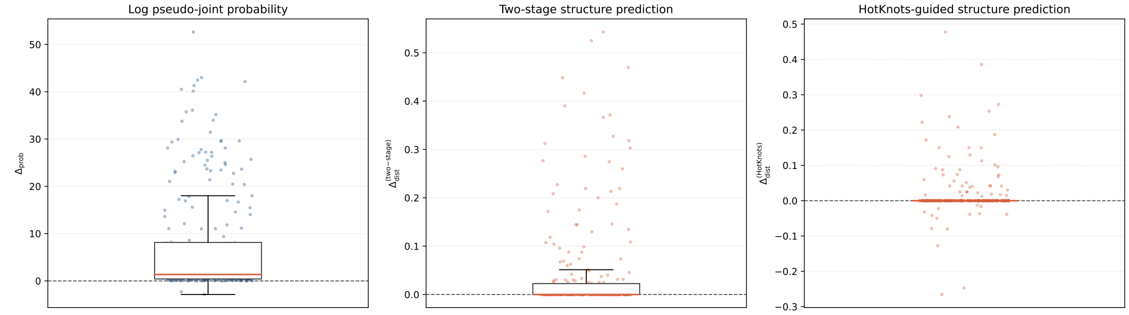


Figure S2: Paired comparison of separate PKProbDesign runs using $G_{\text{big}}$ or $G_{\text{small}}$ as the design scaffold. The left panel shows $\Delta_{\text{prob}}$. All 254 values are finite; $G_{\text{big}}$ is higher for 246 targets and $G_{\text{small}}$ for 8. The middle panel shows $\Delta_{\text{dist}}^{(\text{two-stage})}$; the comparison gives 71 wins for $G_{\text{big}}$, none for $G_{\text{small}}$, and 183 ties. The right panel shows $\Delta_{\text{dist}}^{(\text{HotKnots})}$; the comparison gives 46 wins for $G_{\text{big}}$, 14 for $G_{\text{small}}$, and 194 ties.

### Figure S3

Figure S3 reports the normalized base-pair Hamming-distance summary obtained from IPknot predictions.

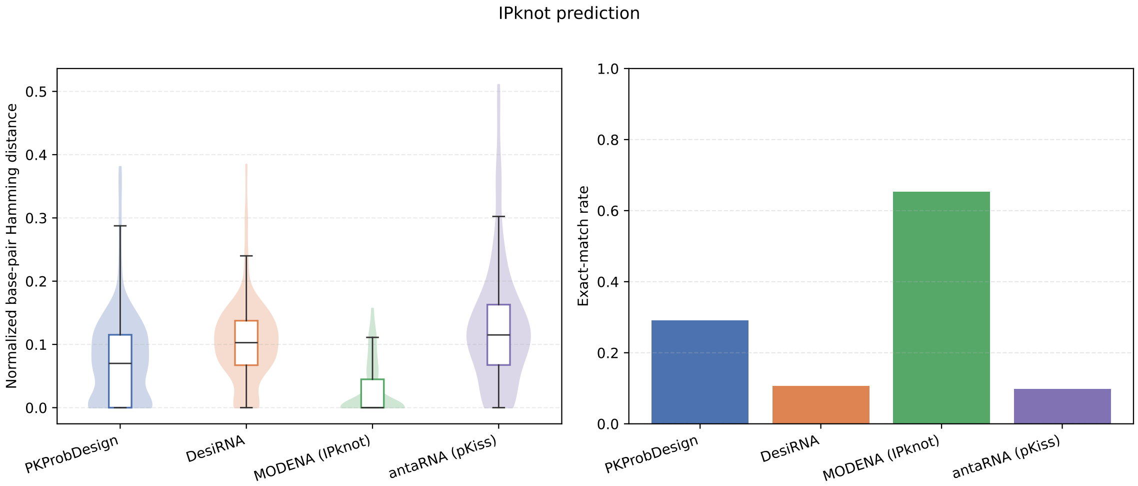


Figure S3: IPknot predictions for the 254 unique density-2 targets. For each method, the representative sequence selected by the $G_{\text{big}}$-scaffold pseudo-joint score was used; the scaffold assignment was not supplied to IPknot. The left panel shows normalized base-pair Hamming distance and the right panel shows exact-match rate. All displayed representative sequences were evaluated with IPknot 1.1.0 using the LinearPartition-C model. Lower distance and higher exact-match rate indicate better agreement with the target structure.

### Text S3

#### Sensitivity to scaffold assignment

For each target, the graph-colouring rule determines $G_{\text{big}}$ and $G_{\text{small}}$ before optimization. The main analysis used $G_{\text{big}}$ as the scaffold, whereas the sensitivity analysis used $G_{\text{small}}$ as the scaffold for every target. PKProbDesign and DesiRNA used this scaffold assignment during sequence design and probability evaluation; MODENA and antaRNA used it only for probability evaluation. For every method, the representative had the highest pseudo-joint score among 20 designs under this assignment; these assignment-specific scores should not be interpreted as two estimates of the same full-structure probability.

With $G_{\text{small}}$ as the scaffold, DesiRNA had the highest pseudo-joint score on 141 targets, PKProbDesign on 110, MODENA on 3, and antaRNA on none (Figure S4). In the paired comparison, PKProbDesign was higher than DesiRNA on 111 targets and lower on 143, as reported in Table S2. DesiRNA was higher for the scaffold term on 233 targets, whereas PKProbDesign was higher for the conditional-extension term on 228. This result is consistent with DesiRNA explicitly optimizing its input scaffold and shows that both probability terms affect the combined score. Across the two scaffold assignments, DesiRNA had a higher assignment-specific score with $G_{\text{big}}$ as the scaffold on 218 targets and with $G_{\text{small}}$ on 36.

On some targets, PKProbDesign had substantially lower scores than DesiRNA when the larger $G_{\text{big}}$ component was used as the extension. Under the fixed optimization budget of 30 samples per iteration for 100 iterations and one fixed seed, the analysis cannot determine whether this reflects limited sampling, a more difficult optimization setting, or both. The corresponding two-stage, HotKnots-guided, and IPknot structure-prediction results are reported in Figures S5 and S6.

### Table S2

Table S2: $G_{\text{small}}$**-scaffold sensitivity analysis.** PKProbDesign and DesiRNA used $G_{\text{small}}$ as the scaffold during sequence design and probability evaluation. The paired difference is defined as PKProbDesign minus DesiRNA. The six-comparison adjustment treats all pairwise method comparisons for one metric as a family; the global adjustment covers all 36 tests in the analysis.

| Metric | $n$ | PKProbDesign better | DesiRNA better | Ties | Median $\Delta$ | Raw $p$ | Six-comparison $p$ | Global $p$ |
|---|---|---|---|---|---|---|---|---|
| Log pseudo-joint folding probability | 254 | 111 | 143 | 0 | $-0.1521$ | $4.50 \times 10^{-8}$ | $2.70 \times 10^{-7}$ | $1.62 \times 10^{-6}$ |
| Log scaffold folding probability | 254 | 21 | 233 | 0 | $-0.5624$ | $3.43 \times 10^{-39}$ | $2.06 \times 10^{-38}$ | $1.23 \times 10^{-37}$ |
| Log conditional extension probability | 254 | 228 | 26 | 0 | 0.6368 | $1.10 \times 10^{-37}$ | $6.62 \times 10^{-37}$ | $3.97 \times 10^{-36}$ |

### Figure S4

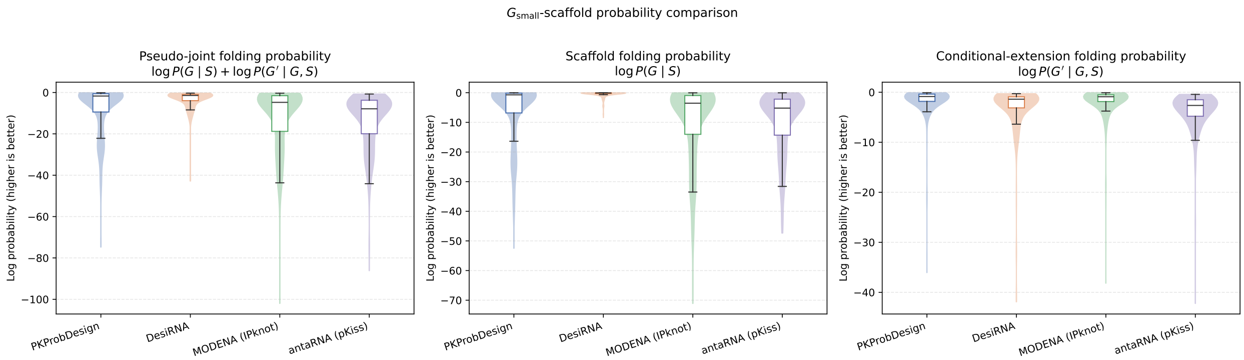


Figure S4: $G_{\text{small}}$-scaffold probability comparison, with $G_{\text{big}}$ as the extension. PKProbDesign and DesiRNA used this scaffold assignment during sequence design and probability evaluation. MODENA and antaRNA did not use this decomposition during design; it was used only to evaluate their sequences and select their representatives. For every target and method, the representative with the highest pseudo-joint score among 20 designs is shown. The left, middle, and right panels show the log pseudo-joint, scaffold, and conditional-extension probabilities, respectively. Higher values indicate higher folding probabilities.

### Figure S5

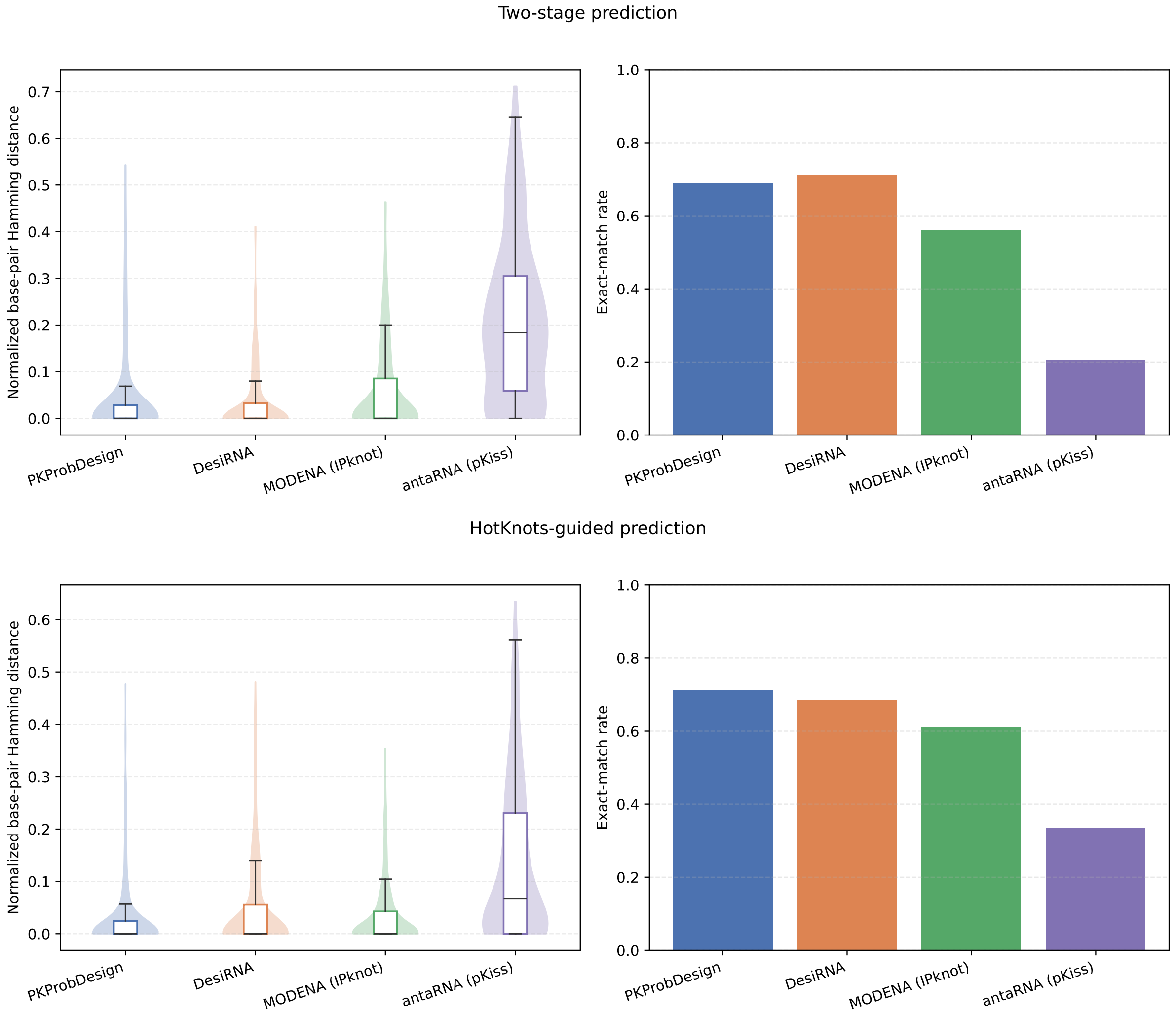


Figure S5: Structure-prediction results for representatives selected by the $G_{\text{small}}$-scaffold pseudo-joint score. The upper row shows two-stage prediction using a LinearFold scaffold; the lower row shows HotKnots-guided prediction. Within each row, the left panel shows normalized base-pair Hamming distance and the right panel shows exact-match rate. The scaffolds used for prediction were obtained from LinearFold or HotKnots, not from the $G_{\text{small}}$ assignment used for probability evaluation. Lower distance and higher exact-match rate indicate better target agreement.

### Figure S6

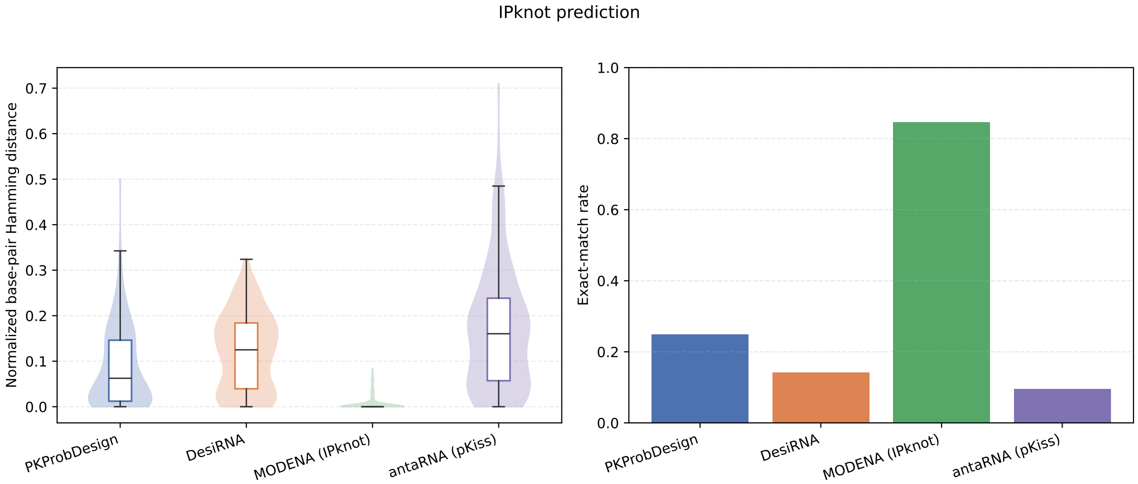


Figure S6: IPknot predictions for representatives selected by the $G_{\text{small}}$-scaffold pseudo-joint score. The assignment is used only for representative selection and is not supplied to IPknot. The left panel shows normalized base-pair Hamming distance and the right panel shows exact-match rate. This evaluation uses the same predictor that MODENA uses during optimization.
